# Transferring and Generalizing Deep-Learning-based Neural Encoding Models across Subjects

**DOI:** 10.1101/171017

**Authors:** Haiguang Wen, Junxing Shi, Wei Chen, Zhongming Liu

## Abstract

Recent studies have shown the value of using deep learning models for mapping and characterizing how the brain represents and organizes information for natural vision. However, modeling the relationship between deep learning models and the brain (or encoding models), requires measuring cortical responses to large and diverse sets of natural visual stimuli from single subjects. This requirement limits prior studies to few subjects, making it difficult to generalize findings across subjects or for a population. In this study, we developed new methods to transfer and generalize encoding models across subjects. To train encoding models specific to a subject, the models trained for other subjects were used as the prior models and were refined efficiently using Bayesian inference with a limited amount of data from the specific subject. To train encoding models for a population, the models were progressively trained and updated with incremental data from different subjects. For the proof of principle, we applied these methods to functional magnetic resonance imaging (fMRI) data from three subjects watching tens of hours of naturalistic videos, while deep residual neural network driven by image recognition was used to model the visual cortical processing. Results demonstrate that the methods developed herein provide an efficient and effective strategy to establish subject-specific or populationwide predictive models of cortical representations of high-dimensional and hierarchical visual features.

## Introduction

An important area in computational neuroscience is developing encoding models to explain brain responses given sensory input (Trappenberg, 2009). In vision, encoding models that account for the complex and nonlinear relationships between natural visual inputs and evoked neural responses can shed light on how the brain organizes and processes visual information through neural circuits (Paninski et al., 2007; Naselaris et al., 2011; Chen et al., 2014; Cox and Dean, 2014; Kriegeskorte, 2015). Existing models may vary in the extent to which they explain brain responses to natural visual stimuli. For example, Gabor filters or their variations explain the neural responses in the primary visual cortex but not much beyond it (Kay et al., 2008; Nishimoto et al., 2011). Visual semantics explain the responses in the ventral temporal cortex but not at lower visual areas (Naselaris et al., 2009; Huth et al., 2012). On the other hand, brain-inspired deep neural networks (DNN) (LeCun et al., 2015), mimic the feedforward computation along the visual hierarchy (Kriegeskorte, 2015; Yamins and DiCarlo, 2016; Kietzmann et al., 2017; van Gerven, 2017), match human performance in image recognition (Krizhevsky et al., 2012; Simonyan and Zisserman, 2014; Szegedy et al., 2015; He et al., 2016), and explain cortical activity over nearly the entire visual cortex in response to natural visual stimuli (Yamins et al., 2014; Güçlü and van Gerven, 2015b, a; Wen et al., 2016, 2017; Eickenberg et al., 2017; Seeliger et al., 2017).

These models also vary in their complexity. In general, a model that explains the brain in natural vision tends to extract a large number of visual features given the diversity of the visual world and the complexity of neural circuits. For DNN, the feature space usually has a very large dimension in the order of millions (Krizhevsky et al., 2012; Simonyan and Zisserman, 2014; Szegedy et al., 2015; He et al., 2016). Even if the model and the brain share the same representations up to linear transform (Yamins and DiCarlo, 2016), matching such millions of features onto billions of neurons or tens of thousands of neuroimaging voxels requires substantial data to sufficiently sample of the feature space and reliably train the linear transformation. For this reason, current studies have focused on only few subjects while training subject-specific encoding models with neural responses observed from each subject given hundreds to thousands of natural pictures (Güçlü and van Gerven, 2015b; Eickenberg et al., 2017; Seeliger et al., 2017), or several to tens of hours of natural videos (Güçlü and van Gerven, 2015a; Wen et al., 2016, 2017; Eickenberg et al., 2017). However, a small subject pool leads to concerns on the generality of the conclusions drawn from such studies. Large data from single subjects are often rare and difficult to collect especially for patients and children. Thus, it is desirable to transfer encoding models across subjects to mitigate the need for a large amount of training data from single subjects, while being able to specifically account for cross-subject differences.

Beyond the level of single subjects, what is also lacking is a method to train encoding models for a group by using data from different subjects in the group. This need rises in the context of “big data”, as data sharing is increasingly expected and executed (Teeters et al., 2008; Van Essen et al., 2013; Paltoo et al., 2014; Poldrack and Gorgolewski, 2014). For a group of subjects, combining data across subjects can yield much more training data than are attainable from a single subject. A population-wise encoding model also sets the baseline for identifying any individualized difference within a population. However, training such models with a very large and growing dataset as a whole is computationally inefficient or even intractable with the computing facilities available to most researchers (Fan et al., 2014).

Here, we developed methods to train DNN-based encoding models for single subjects or multiple subjects as a group. Our aims were to 1) mitigate the need for a large training dataset for each subject, and 2) efficiently train models with big and growing data combined across subjects. To achieve the first aim, we used pre-trained encoding models as the prior models in a new subject, reducing the demand for collecting extensive data from the subject in order to train the subject-specific models. To achieve the second aim, we used online learning algorithm (Fontenla-Romero et al., 2013) to adjust an existing encoding model with new data to avoid retraining the model from scratch with the whole dataset. To further leverage both strategies, we employed functional hyperalignment (Guntupalli et al., 2016) between subjects before transferring encoding models across subjects. Using experimental data for testing, we showed the merits of these methods in training the DNN-based encoding models to predict functional magnetic resonance imaging (fMRI) responses to natural movie stimuli in both individual and group levels.

## Methods and Materials

### Experimental data

In this study, we used the video-fMRI data from our previous studies (Wen et al., 2016, 2017). The fMRI data were acquired from three human subjects (Subject JY, XL, and XF, all female) when watching natural videos. The videos covered diverse visual content representative of real-life visual experience.

For each subject, the video-fMRI data was split into three independent datasets for 1) functional alignment between subjects, 2) training the encoding models, and 3) testing the trained models. The corresponding videos used for each of the above purposes were combined and referred to as the “alignment” movie, the “training” movie, and the “testing” movie, respectively. For Subjects XL and XF, the alignment movie was 16 minutes; the training movie was 2.13 hours; the testing movie was 40 minutes. To each subject, the alignment and training movies were presented twice, and the testing movie was presented ten times. For Subject JY, all the movies for Subjects XL and XF were used; in addition, the training movie also included 10.4 hours of new videos not seen by Subjects XL and XF, which were presented only once.

Despite their different purposes, these movies were all split into 8-min segments, each of which was used as continuous visual stimuli during one session of fMRI acquisition. The stimuli (20.3^o^ × 20.3^o^) were delivered via a binocular goggle in a 3-T MRI system. The fMRI data were acquired with 3.5 mm isotropic resolution and 2 s repetition time, while subjects were watching the movie with eyes fixating at a central cross. Structural MRI data with T_1_ and T_2_ weighted contrast were also acquired with 1 mm isotropic resolution for every subject. The fMRI data were preprocessed and coregistered onto a standard cortical surface template (Glasser et al., 2013). More details about the stimuli, data acquisition and preprocessing are described in our previous papers (Wen et al., 2016, 2017).

### Nonlinear feature model based on deep neural network

The encoding models took visual stimuli as the input, and output the stimulus-evoked cortical responses. As shown in Fig. 1, it included two steps. The first step was a nonlinear feature model, converting the visual input to its feature representations; the second step was a voxel-wise linear response model, projecting the feature representations onto the response at each fMRI voxel (Kay et al., 2008; Naselaris et al., 2009; Nishimoto et al., 2011; Huth et al., 2012; Güçlü and van Gerven, 2015b, a; Wen et al., 2016, 2017; Eickenberg et al., 2017; Seeliger et al., 2017). The feature model is described in this subsection, and the response model is described in the next sub-section.

**Figure 1.**
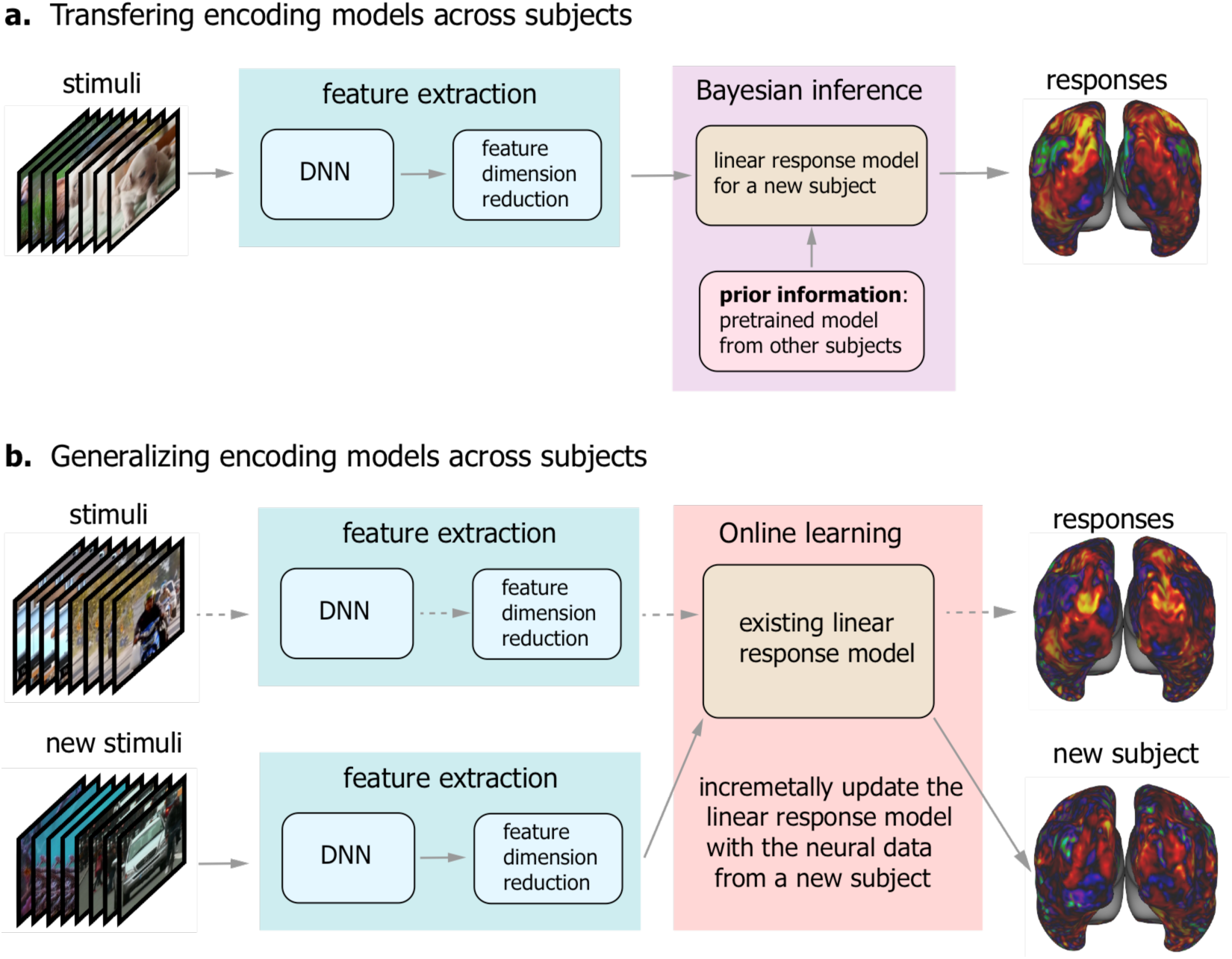
Schemes of transferring and generalizing DNN-based neural encoding models across subjects. (a) Transferring encoding models across subjects. The encoding model comprises the nonlinear feature model and the linear response model. In the feature model, the feature representation is extract from the visual stimuli through the deep neural network (DNN), and followed by the feature dimension reduction. In the response model, the model parameters are estimated by using Bayesian inference with subject-specific neural data as well as a prior model trained from other subjects. (b) Generalizing encoding models across subjects. The dash arrows indicate the existing encoding model trained with the data from a group of subjects. The existing model can be incrementally updated by using the new data from a new subject with an online learning algorithm. In the scheme, the feature model is common any subjects and any stimuli, and the response model will be updated when new subject data is available.

In line with previous studies (Güçlü and van Gerven, 2015b, a; Wen et al., 2016, 2017; Eickenberg et al., 2017; Seeliger et al., 2017), a deep neural network (DNN) was used in the present study as the feature model to extract hierarchical features from visual input. Here, a specific version of the DNN, i.e. deep residual network (ResNet) (He et al., 2016), was used for this purpose. Briefly, the ResNet was composed of 50 hidden layers of nonlinear computational units that encoded increasingly abstract and complex visual features (He et al., 2016). Passing an image into the ResNet yielded an activation value at each unit. Passing every frame of a movie into the ResNet yielded an activation time series at each unit, indicating the time-varying representation of a specific feature in the movie. In this way, the feature representations of the training and testing movies could be extracted, as in previous studies (Wen et al., 2016, 2017).

### Feature dimension reduction

The feature space in the ResNet had a huge dimension over 10^6^. This dimensionality could be reduced since individual features were not independent. For this purpose, principal component analysis (PCA) was applied first to each layer and then across layers. To define a set of principal components generalizable across various visual stimuli, a training movie as long as 12.54 hours was used to sample the original feature space. The corresponding feature representations were demeaned and divided by its standard deviation, yielding the standardized feature representation at each unit. Then, PCA was applied to the standardized feature representations from all units in each layer, as expressed as Eq. (1).

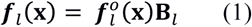

where
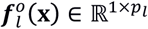 stands for the standardized feature representation from layer *l* given a visual input **x**, **B**_*l*_ ∈ ℝ^*p_l_*×*q_l_*^ consists of the principal components (unit column vectors) for layer *l*, ***f*(x)** ∈ ℝ^1×*ql*^ is the feature representation after reducing the dimension from to *p_l_* to *q_l_*.

Due to the high dimensionality of the original feature space and the large number of video frames, we used an efficient singular value decomposition updating algorithm (or SVD-updating algorithm) (Zha and Simon, 1999; Zhao et al., 2006) to obtain the principal components *B*_*l*_. Briefly, the 12.54-hour training movie was divided into blocks, where each block was defined as an 8-min segment (i.e. a single fMRI session). The principal components of the feature representation were initially calculated from a block and were incrementally updated with new blocks by keeping >99% variance of the feature representations of every block of visual input **x** (see **SVD-updating algorithm in Supplementary Information** for details).

Following the layer-wise dimension reduction, PCA was applied to the feature representations from all layers with SVD-updating algorithm, keeping the principal components that explained >99% variance across layers for every block of visual stimuli. The final dimension reduction was implemented as Eq. (2).

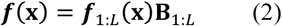

where
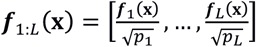 stands for the feature representation concatenated across *L* layers, **B**_1:*L*_ consists of the principal components of ***f***_1:*L*_(**x**) given the 12.54-hour training movie, and ***f***(**x**) ∈ ℝ^l×*k*^ is the final dimension-reduced feature representation.

The principal components **B**_*l*_ and **B**_1:*L*_ together defined a dimension-reduced feature space, and their transpose defined the transformation to the original feature space. So, given any visual stimulus **x**, its dimension-reduced feature representation could be obtained through Eqs. (1) and (2) with fixed **B**_*l*_ and **B**_1:*L*_. Once trained, the feature model including the feature dimension reduction, was assumed to be common to any subjects and any stimuli.

### Voxel-wise linear response model

As the second part of the encoding model, a voxel-wise linear regression model was trained to predict the response *r_v_* (**x**) at voxel *v* evoked by the stimulus **x** based on the dimension-reduced feature representation of the stimulus, ***f*(x)**, as expressed by Eq. (3).

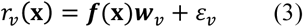

where ***w_v_*** is a column vector of unknown regression coefficients specific to voxel *v*, and *ε_v_* is the noise (unexplained by the model). Here, the noise was assumed to follow a Gaussian distribution with zero mean and variance equal to 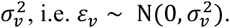

Eq. (3) can be rewritten in vector/matrix notations as Eq. (4) for a finite set of visual stimuli (e.g. movie frames).

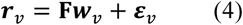

where **F** ∈ ℝ^*n*×*k*^ stands for the feature representations of *n* stimuli, ***r_v_*** ∈ ℝ^*n*×l^ is the corresponding evoked responses, and 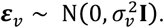

To estimate the regression coefficients ***w_v_*** in Eq. (4), we used and compared two methods: 1) maximum likelihood estimation (MLE) (Paninski, 2004; Ahrens et al., 2008; Trappenberg, 2009), by maximizing the probability of the observed response *r_v_* (**x**) given the stimulus **x**; 2) Bayesian inference (or maximum a posteriori estimation) (Sahani and Linden, 2003; Paninski et al., 2007), by maximizing the posterior probability of ***w**_v_* given the stimulus **x**, the observed fMRI response *r_v_*(**X**), and the prior knowledge about ***w**_v_*. The former was used for training subject-specific encoding models with subject-specific training data; the latter was what we proposed herein for transferring encoding models across subjects.

### Training the response model with maximum likelihood estimation

From Eq. (4), the likelihood of the response ***r**_v_* given the unknown coefficients ***w**_v_* and the known feature representations **F** followed a multivariate Gaussian distribution, as Eq. (5).

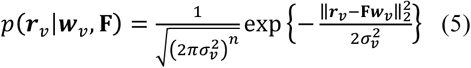

The MLE of ***w**_v_* was obtained by maximizing the log of this likelihood, which was equivalent to minimizing the objective function as Eq. (6), with additional regularization penalty to avoid overfitting (Banerjee et al., 2014).

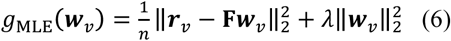

where the first term is the mean square error, the second term is the L2-regularization penalty, and *λ* is the regularization parameter. The analytical solution to minimizing (6) is as Eq. (7)

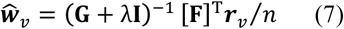

where **G** = [**F**]^T^**F**/*n* is the covariance matrix of **F**.

### Training the response model with Bayesian inference with a model prior

When prior knowledge about ***w**_v_* was available, ***w**_v_* could be estimated with Bayesian inference. In this regard, ***w**_v_* was modeled as a multivariate random variable. In this study, the prior knowledge about ***w**_v_* was its expectation, 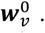 Assuming that ***w**_v_* followed a multivariate Gaussian distribution with its covariance 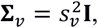 the probability of ***w**_v_* was thus written as Eq. (8).

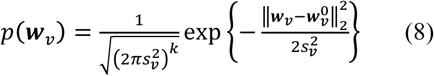

Note that this prior distribution was independent of any particular visual stimuli and thus their feature representations, i.e. *p*(***w**_v_*) = *p*(***w**_v_* |**F**). Therefore, given **F** and ***r**_v_*, the posterior distribution of ***w**_v_* was as Eq. (9) according to the Bayes’ rule.

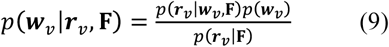

where *p*(***r_v_*|F**) was constant since ***r**_v_* and **F** were known.

Combining Eqs. (5), (8) and (9), maximizing the posterior probability of ***w**_v_* was equivalent to minimizing the following objective function with additional L_2_-regularization to avoid overfitting.

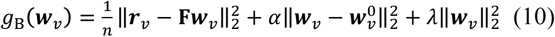

where 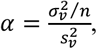 and the last term is the L_2_-regularization penalty. Note that if 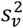 = + ∞, i.e. *α* = 0, the prior is non-informative, and the objective function in Eq. (10) becomes equivalent to that in Eq. (6). The analytical solution to minimizing (10) was as Eq. (11).

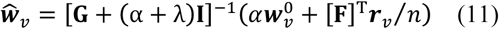

where **G** = [**F**]^T^**F**/*n* is the covariance matrix of **F**.

### Choosing hyper-parameters with cross-validation

The hyper-parameters *λ* in Eq. (7) or (*α*, *λ*) in Eq. (11) were determined for each voxel by four-fold cross-validation (Geisser, 1993). Specifically, the training video-fMRI dataset was divided into four subsets of equal size: three for the model estimation, and one for the model validation. The validation accuracy was measured as the correlation between the predicted and measured cortical responses. The validation was repeated four times such that each subset was used once for validation. The validation accuracy was averaged across four repeats. Finally, the hyper-parameters were chosen such that the average validation accuracy was maximal.

### Testing the encoding performance with the testing movie

Once voxel-wise encoding models were trained with either the MLE or Bayesian methods, we evaluated the model performance in predicting the cortical responses to the testing movie that was never used for training the encoding model (either the feature model or the response model). The prediction accuracy was quantified as the correlation (*r*) between the predicted and observed fMRI responses at each voxel given the testing movie. Since the testing movie included five different 8-min sessions with entirely different movie content, the prediction accuracy was evaluated separately for each session and then averaged across sessions. The significance of the average voxel-wise prediction accuracy was evaluated with a block-permutation test (Adolf et al., 2014) with a block length of 30 seconds (corrected at false discovery rate (FDR) *q* < 0.01), as used in our prior study (Wen et al., 2016, 2017).

### Evaluating the MLE-based encoding models

When MLE was used to train the linear response model for each voxel, the model parameters were derived without any prior information but entirely based on the training data. Here, we tested how the model performance evaluated with the testing movie depended on the size of the training data.

To do so, we trained the encoding models for Subject JY using a varying part of the 10.4-hour training data. The data used for model training ranged from 16 minutes to 10.4 hours. For such models trained with varying lengths of data, we tested their individual performance in predicting the responses to the 40-min testing movie. We calculated the percentage of predictable voxels (i.e. significant with the block-permutation test) out of the total number of cortical voxels, and evaluated it as a function of the size of the training data. We also evaluated the histogram of the prediction accuracy for all predictable voxels, and calculated the overall prediction accuracy in regions of interest (ROIs) (Glasser et al., 2016) by averaging across voxels within ROIs.

### Evaluating the Bayesian-estimated encoding models

When Bayesian inference was used to train the linear response model with a prior knowledge about the model, the model parameters depended on both the model prior and the new training data. As such, one might not require so many data to train the model as required without any prior. We used this Bayesian method for transferring encoding models from one subject to another.

Specifically, we used MLE to train the models based on the 10.4-hour training data from one subject (JY), and used the trained models as the model prior for other subjects (XF and XL). With this prior model from Subject JY, we used the Bayesian method to train the encoding models for Subject XF and XL based on either short (16 minutes, i.e. two 8-min sessions) or long (2.13 hours, i.e. 16 sessions) training data specific to them. Note that the movie used for training the prior model in Subject JY was different from either the training or testing movies for Subject XL and XF. With either short or long training data, we evaluated the model performance in predicting the responses to the testing movie for Subject XF and XL. We compared such trained models with those trained by using MLE with the same training data but without any prior, or the prior models without retraining with any data from Subject XF and XL.

The comparison was made with respect to the number of predictable voxels and the voxel-wise prediction accuracy (after converting the correlation coefficients to the z scores with the Fisher’s r-to-z transform, i.e. z = arctanh(*r*)). Such performance difference between models was evaluated when different two sessions of the training movie were used for training and when different sessions of the testing movie were used for testing. The resulting difference was further tested for significance with one-sample t-test (corrected at false discovery rate (FDR)*q*<0.01).

### Hyperalignment between subjects

We also explored whether transferring encoding models from one subject to another would also benefit from performing functional hyperalignment before using the Bayesian algorithm. Specifically, we used the searchlight hyperalignment algorithm (Guntupalli et al., 2016) to account for the individual difference in fine-scale functional architecture beyond anatomical alignment (Glasser et al., 2013). Given the 16-min alignment movie, the fMRI responses within a searchlight (with a radius of 20mm) were viewed as a high-dimensional vector that varied in time. A Procrustes transformation (Schönemann, 1966) was optimized to align high-dimensional response patterns between subjects (Guntupalli et al., 2016).

To evaluate the effect of hyperalignment in transferring encoding models across subjects, we applied the searchlight hyperalignment from Subject JY to Subject XL and XF. After hyperalignment, we used the Bayesian method to train encoding models for Subject XL and XF, by incorporating the prior models as the encoding models for the source subject (Subject JY) functionally aligned to the target subject (Subject XL or XF). The performance (i.e. predicting the response to the testing movie) of such encoding models trained with hyperalignment was evaluated and compared with those without hyperalignment. The comparison was made with respect to the number of predictable voxels and the voxel-wise prediction accuracy. The resulting difference was evaluated and tested for significance with one-sample t-test corrected at false discovery rate (FDR) *q*<0.01.

### Training group-level encoding models with online learning

Here, we describe an online learning algorithm (Fontenla-Romero et al., 2013) to train group-level encoding models based on different video-fMRI data acquired from different subjects, by extending the concept of online implementation for the Levenberg-Marquardt algorithm (Dias et al., 2004). The central idea is to update the encoding models trained with existing data based on the data that become newly available.

Suppose that existing training data are available for a set of visual stimuli, **X**^0^ (*n*^0^ samples). Let **F**^0^ be the corresponding feature representations after dimension reduction, 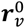 be the responses at voxel *v*. Let 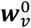 be the regression parameters in the voxel-specific encoding models trained with **F**^0^ and 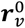 according to Eq. (7). Given incremental training data, **X**^1^ (*n*^1^ samples), **F**^1^ and 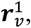 the parameters in the updated encoding model can be obtained by minimizing the objective function below.

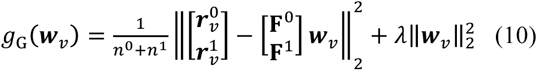

The optimal solution is expressed as Eq. (11).

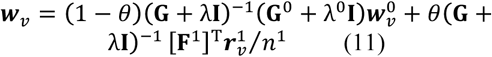

where **G**^0^ = [**F**^0^]^T^**F**^0^/*n*^0^ and **G**^1^ = [**F**^1^]^T^**F**^1^/*n*^1^ are the covariance matrices of **F**^0^ and **F**^1^, respectively; **G** = (1 – *θ*)**G**^0^ + *θ***G**^1^ is their weighted sum where the parameter *θ* specifies the relative weighting of the new data and the previous data. See **Derivation of group-level encoding models** in **Supplementary Information** for the derivation of Eq. (11). In this study, *θ* was set as the ratio of the corresponding sample sizes, i.e. 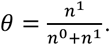 As such, the samples in the new data were assumed to be as important as those in the previous data.

According to Eq. (11), the encoding model could be incrementally updated by incorporating new data without training the model from scratch. See **Algorithm 1** in **Supplementary Information** for the updating rules. As more and more data was used for model training, the encoding model was expected to converge, as (**G** + λ**I**)^1^(**G**^0^ + λ^0^**I** → **I** and *θ* → 0. When it was used to utilize the growing training data from different subjects, this algorithm converged to the group-level encoding models.

As a proof of concept, we trained group-level encoding models by incrementally updating the models with 16-min video-fMRI training data sampled from each of the three subjects in the group. Before each update, the incremental fMRI data was functionally aligned to the data already used to train the existing models (see ***Hyperalignment between subjects*** in ***Methods***). After the encoding models were trained with all the training data combined across all the subjects, we evaluated their prediction performance given the testing movie for each subject. The prediction accuracy of the group-level encoding models was averaged across subjects. We then compared the prediction performance before and after every update by incorporating new data.

## Results

In recent studies, DNNs driven by image or action recognition were shown to be able to model and predict cortical responses to natural picture or video stimuli (Khaligh-Razavi and Kriegeskorte, 2014; Yamins et al., 2014; Güçlü and van Gerven, 2015b, a; Cichy et al., 2016; Wen et al., 2016, 2017; Eickenberg et al., 2017; Seeliger et al., 2017). This ability rested upon encoding models, in which non-linear features were extracted from visual stimuli through DNNs and the extracted features were projected onto stimulus-evoked responses at individual locations through linear regression. Herein, we investigated the amount of data needed to train DNN-based encoding models in individual subjects, and developed new methods for transferring and generalizing encoding models across subjects without requiring extensive data from single subjects.

### Encoding performance depended on the size of the training data

In this study, we focused on a specific DNN (i.e. ResNet) – a feed-forward convolutional neural network (CNN) pre-trained for image recognition (He et al., 2016). The ResNet included 50 successive layers of computational units, extracting around 10^6^ non-linear visual features. This huge dimensionality could be reduced by two orders of magnitude, by applying PCA first to every layer and then across all layers (see ***Feature dimension reduction*** in ***Methods***). The reduced feature representations were able to capture 99% of the variance of the original features in every layer.

Despite the reduction of the feature dimensionality, training a linear regression model to project the feature representations onto the fMRI response at each voxel still required a large amount of data if the model was estimated solely based on the training data with the MLE method (***Training the response model with maximum likelihood estimation*** in ***Methods***). Here, we evaluated the effects of the size of the training data on the performance of the MLE-estimated encoding models in terms of predicting the responses to the testing movie, of which the data were not used for training to ensure unbiased testing. When trained with 10.4 hours of video-fMRI data, the prediction accuracy of the encoding models was statistically significant (permutation test, FDR q<0.01) for nearly the entire visual cortex (Fig. 2.a). The number of predictable voxels and the prediction accuracy were notably reduced as the training data were reduced to 5.87 hours, 2.13 hours, or 16 minutes (Fig. 2.b). With increasing sizes of training data, the predictable areas increased monotonically, rising from about 20% of the cortical surface with 16-min of training data, while reaching >40% of the cortical surface with 10.4-hour of data (Fig. 2.c). The average prediction accuracies, although varying across regions of interest (ROIs), showed an increasing trend as a growing amount of data were used for model training (Fig. 2.d). It appeared that the trend did not stop at 10.4 hours, suggesting a sub-optimal encoding model even if trained with such a large set of training data. Therefore, using MLE to train encoding models for a single subject would require at least 10 hours of video-fMRI data from the same subject.

**Figure 2.**
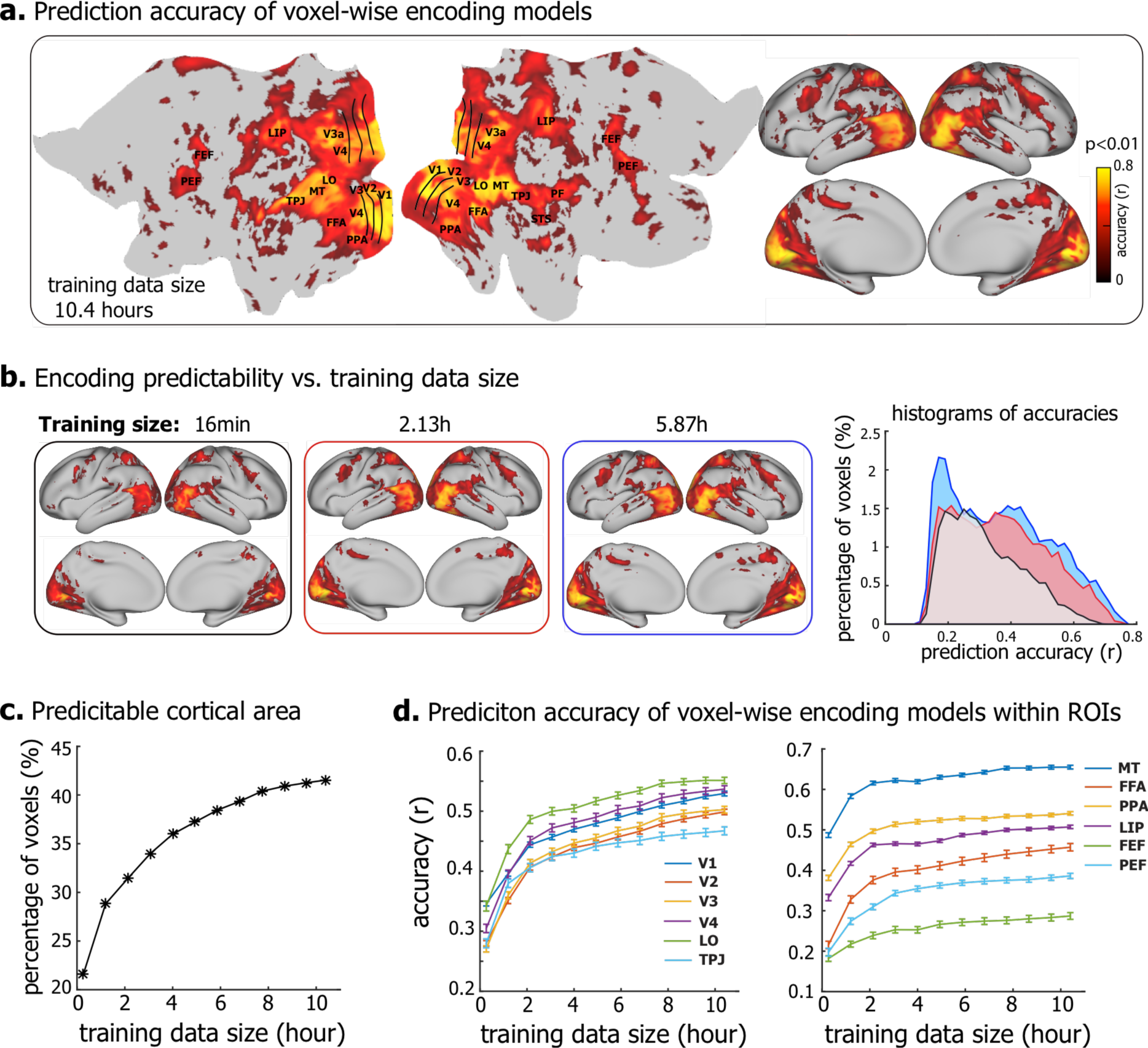
DNN-based neural encoding models for Subject JY. **(a)** Performance of neural encoding models (trained with 10.4-hour data) in predicting the cortical responses to novel testing movies. The accuracy is measured by the average Pearson’s correlation coefficient (r) between the predicted and the observed fMRI responses across five testing movies (permutation test, q<0.01 after correction for multiple testing using the false discovery rate (FDR) method). The prediction accuracy is visualized on both flat (left) and inflated (right) cortical surfaces. **(b)** Prediction accuracy of encoding models trained with less training data, i.e. 16min, 2.13h, and 5.87h. The right is the histograms of prediction accuracies. The x-axis is the prediction accuracy ranging from 0 to 0.8, divided into bins of length *Δr*=0.02, the y-axis is the percentages of predictable voxels in the cortex within accuracy bins. **(c)** The percentage of predictable voxels as a function of training data size ranging from 16min to 10.4 hours. **(d)** ROI-level prediction accuracies as functions of the training data size. The error bar indicates the standard error across voxels.

### Transferring encoding models across subjects through Bayesian inference

To mitigate this need for large training data from every subject, we asked whether the encoding models already trained with a large amount of training data could be utilized as the prior information for training the encoding models in a new subject with much less training data. To address this question, we used the encoding models trained with 10.4 hours of training data from Subject JY as *a priori* models for Subject XF and XL. A Bayesian inference method was used to utilize such prior models for training the encoding models for Subject XF and XL with either 16-min or 2.13-hour training data from these two subjects (see ***Training the response model with Bayesian inference with a model prior*** in **Methods**). The resulting encoding models were compared with those trained by using the MLE method (without using any prior models) with the same amount of training data in terms of their accuracies in predicting the responses to the testing movie.

Fig. 3 shows the results for the model comparison in Subject XF. When the training data were as limited as 16 minutes, the encoding models trained with the Bayesian method significantly outperformed those trained with the MLE method (Fig. 3.a). The cortical areas predictable by the Bayesian-estimated encoding models were 26% of the entire cortex, nearly twice as large as those predictable by the MLE-estimated models (14.9% of the entire cortex). Within the predictable areas, the prediction accuracy was also significantly higher with the Bayesian method than with the MLE method (Δ*z* = 0.155 ± 0.0006, one-sample t-test, p<10^−5^) (Fig.3.a). The difference in voxel-wise prediction accuracy was significant (one-sample t-test, p<0.01) in most of the visual areas, especially for those in the ventral stream (Fig. 3.a). The advantage of using the prior model with the Bayesian method largely diminished when 2.13-hour training data were used for training the encoding models (Fig. 3.b). Although larger training data improved the model performance, the improvement was much more notable for the MLE method than for the Bayesian method. For the former, the predictable area increased from 14.9% to 26.7% (of the cortex)(p=6.5× 10^−5^, paired t-test); for the latter, it increased from 26.0% to 28.5% (p=0.017, paired t-test). The differences in the predictable area and the prediction accuracy were relatively minor, although observable (Fig. 3.b). Consistent results were observed in Subject XL (Supplementary Fig. S1). It was noteworthy that the prediction accuracy of the Bayesian-estimated encoding model with 16-min fMRI data was comparable to the MLE-estimated models with 2.13-hour fMRI data (Fig. 3 and Supplementary Fig. S2).

**Figure 3.**
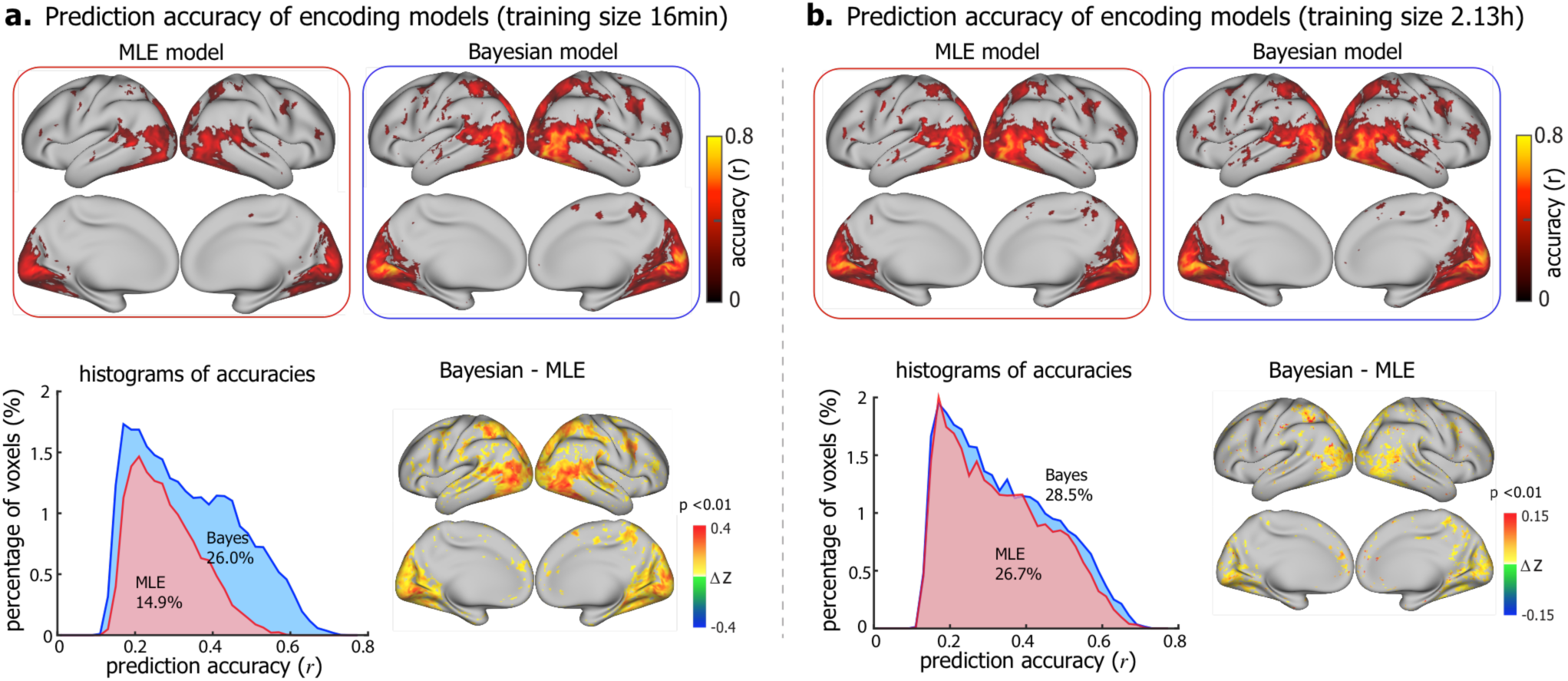
Comparison between Bayesian encoding models and MLE encoding models. Voxel-wise prediction accuracy of encoding models trained with 16min (a) and 2.13h (b) video-fMRI data (permutation test, corrected at FDR q<0.01). The top shows the voxel-wise prediction accuracy of Bayesian (right) and MLE (left) encoding models. The bottom left is the histograms of prediction accuracies of Bayesian (blue) and MLE (red) encoding models. The colored numbers are the total percentages of predictable voxels. The bottom right is the difference of prediction accuracy (Fisher’s z-transformation of r, i.e. z = arctanh(*r*)) between Bayesian and MLE models (one-sample t-test, p<0.01). The figure shows the results for Subject XF, see Figure S1 for Subject XL.

We also asked whether the better performance of the Bayesian-estimated encoding models was entirely attributable to the prior models from a different subject, or it could be in part attributable to the information in subject-specific training data. To address this question, we directly used the prior models (trained with data from Subject JY) to predict the cortical responses to the testing movie in Subject XL and XF. Even without any further training, the prior models themselves yielded high prediction accuracy for widespread cortical areas in Subject XF for whom the models were not trained (Fig. 4.a). When the prior models were fine-tuned with a limited amount (16-min) of training data from this specific subject, the encoding performance was further improved (Fig. 4.b). The improvement was more notable when more (2.13-hour) training data were utilized for refining the encoding models (Fig. 4.c). Similar results were also observed in another subject (Supplementary Fig. S3). Hence, using the Bayesian inference to transfer encoding models across subjects helped to train encoding models for new subjects without requiring extensive training data from them. The subject-specific training data served to tailor the encoding models from the source subject towards the target subject.

**Figure 4.**
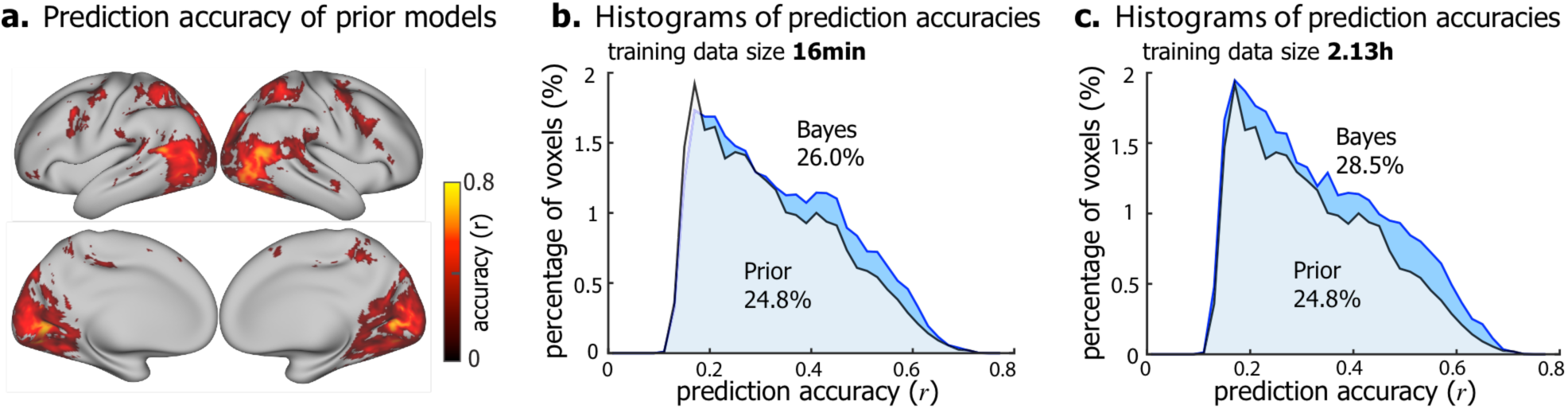
Comparison between Bayesian encoding models and prior encoding models. (a) Voxel-wise prediction accuracy by directly using the prior encoding models (from Subject JY) to predict the responses to novel testing movies for Subject XF (permutation test, corrected at FDR q<0.01). (b) and (c) show the histograms of prediction accuracies of Bayesian (blue) and prior (green) encoding models trained with 16min (b) and 2.13h (c) training data, respectively. See Figure S3 for Subject XL.

### Functional alignment better accounted for individual differences

Transferring encoding models across subjects were based on the assumption that the models and data from individual subjects were co-registered. Typically, the co-registration was based on anatomical features (i.e. anatomical alignment) (Glasser et al., 2013). We expected that searchlight hyperalignment of multi-voxel responses could better co-register the cortical representational space between subjects (Guntupalli et al., 2016) to improve the efficacy of transferring the encoding models across subjects (see ***Hyperalignment between subjects*** in **Methods**).

Therefore, we performed searchlight hyperalignment such that Subject JY’s fMRI responses to the alignment movie were aligned to the other subjects’ responses to the same movie. After applying the same alignment to the encoding models trained for Subject JY, we used the aligned encoding models as the prior model for training the encoding models for Subject XF or XL with 16-min training datasets from each of them. It turned out that using the functional alignment as a preprocessing step further improved the performance of the Bayesian-estimated encoding models. For Subject XF, the model-predictable areas increased from 26% to 27.8% (p=9.7×10^−4^, paired t-test), and the prediction accuracy also increased, especially for the extrastriate visual areas (Fig. 5). Similar results were obtained for Subject XL (Supplementary Fig. S4).

**Figure 5.**
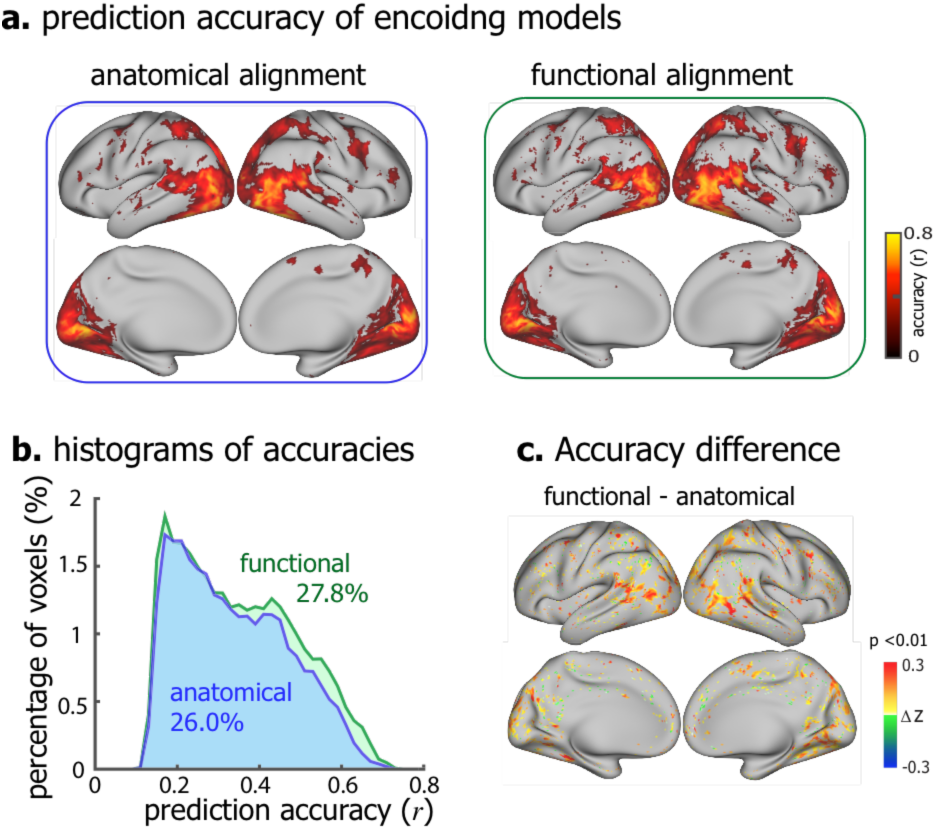
Comparison of Bayesian encoding models between anatomical alignment and functional alignment. (a)Voxel-wise prediction accuracy of Bayesian encoding models based on anatomical alignment (left) and functional alignment (right) (permutation test, corrected at FDR q<0.01). (b) The histograms of prediction accuracies of anatomically aligned (blue) and functionally aligned (green) Bayesian encoding models. The colored numbers are the total percentages of predictable voxels. (c) The voxel-wise difference in prediction accuracy (Fisher’s z-transformation of r, i.e. z=arctanh(*r*)) between functional alignment and anatomical alignment (one-sample t-test, p<0.01). The figure shows the results for Subject XF, see Figure S5 for Subject XL.

### Group-level encoding models

We further explored and tested an online learning strategy to train general encoding models for a group of subjects by incrementally using data from different subjects for model training (see ***Training group-level encoding models with online learning*** in ***Methods***).

Basically, incremental neural data (16 minutes) was obtained from a new subject with new visual stimuli, and was used to update the existing encoding models (Fig. 6a). Such learning strategy allowed training group-level encoding models. The models significantly predicted the cortical response to novel testing movie for each subject (Fig. 6b). With every incremental update, the encoding models predicted wider cortical area covering from 18.4% to 21.72% to 24.27% of the cortex, and achieved higher prediction accuracy within the predictable area (first update: Δ*z* = 0.05 ± 0.0006, p<10^−5^; second update: Δ*z* = 0.036 ± 0.00034, p<10^−5^, one-sample t-test) (Fig. 6.b). Meanwhile, the group-level encoding models exhibited similar predictability across individual subjects (Supplementary Fig. S5).

**Figure 6.**
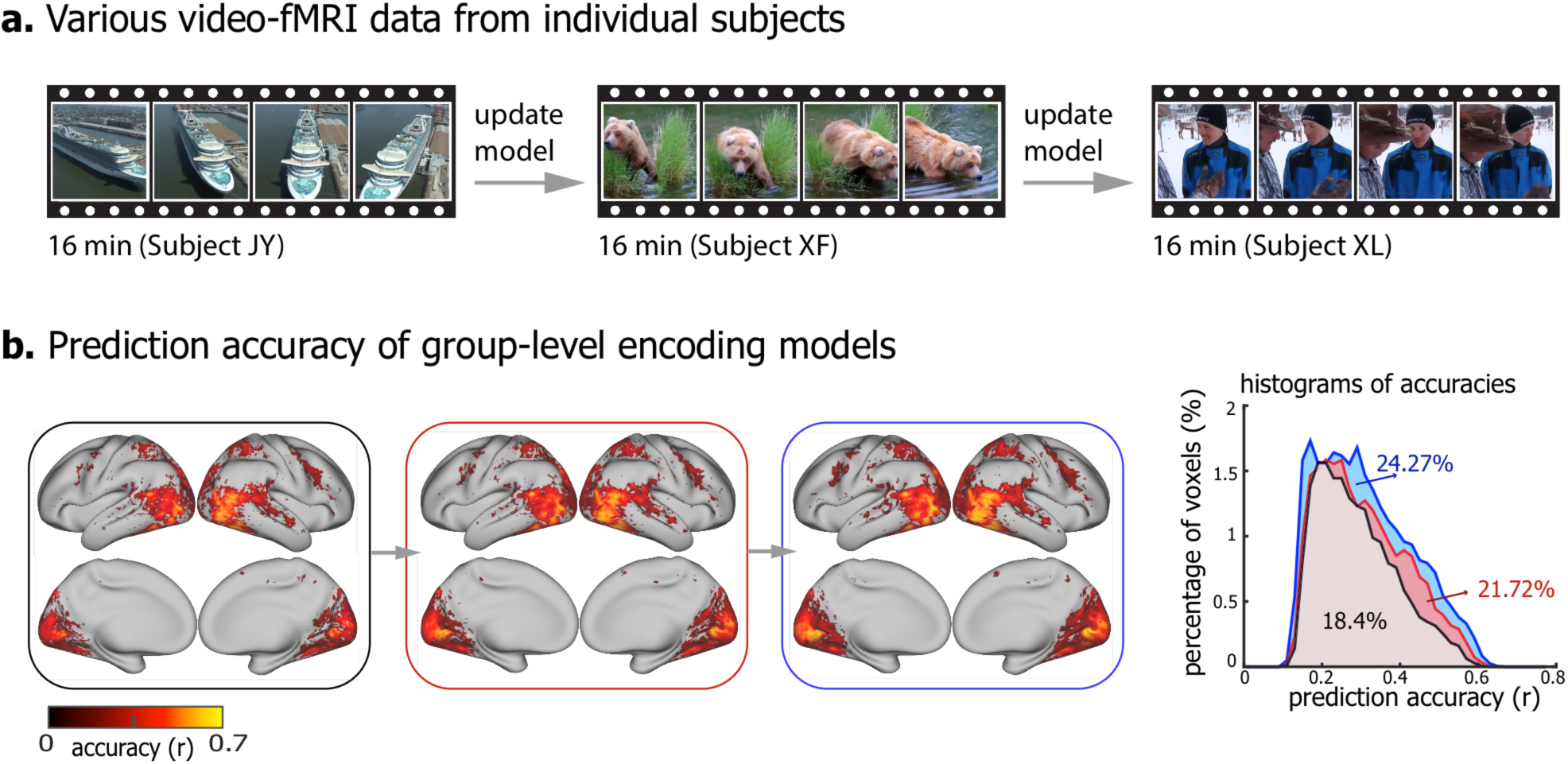
Group-level encoding models. (a) Distinct video-fMRI dataset obtained from different subjects when watching different natural videos. (b) The voxel-wise prediction accuracy of group-level encoding models before and after every incremental update (permutation test, corrected at FDR q<0.01). The right is the histograms of prediction accuracies of incrementally updated encoding models. The colored numbers are the total percentages of predictable voxels. The testing accuracy is averaged across three subjects.

## Discussion

In this article, we have described methods to transfer and generalize encoding models of natural vision across human subjects. Central to our methods is the idea of taking the models learnt from data from one subject (or a group of subjects) as the prior models for training the models for a new subject (or a new group of subjects). This idea, implemented in the framework of Bayesian inference, allows to train subject-specific encoding models with a much less amount of training data than otherwise required if training the models from scratch without considering any model prior. The efficacy of this method, as demonstrated in this paper, suggests that different subjects share largely similar cortical representations of vision (Hasson et al., 2004; Haxby et al., 2011; Conroy et al., 2013; Chen et al., 2017). It has also led us to develop a method to train encoding models generalizable for a population by incrementally learning from different training data collected from different subjects.

The methods are described in the context of using DNN as a feature model, but are also valuable and applicable to other models of visual or conceptual features (Kay et al., 2008; Naselaris et al., 2009; Nishimoto et al., 2011; Huth et al., 2012). In general, the larger the feature space is, the more data is required for training the model that relates the features to brain responses in natural vision. DNNs attempt to extract hierarchical visual features in many levels of complexity (Krizhevsky et al., 2012; Simonyan and Zisserman, 2014; Zeiler and Fergus, 2014; LeCun et al., 2015; Szegedy et al., 2015; He et al., 2016), and thus it is so-far most data demanding to model their relationships to the visual cortex. Nevertheless, DNNs are of increasing interest for natural vision (Cox and Dean, 2014; Khaligh-Razavi and Kriegeskorte, 2014; Yamins et al., 2014; Güçlü and van Gerven, 2015b, a; Kriegeskorte, 2015; Cichy et al., 2016; Wen et al., 2016, 2017; Eickenberg et al., 2017; Horikawa and Kamitani, 2017; Seeliger et al., 2017). Recent studies have shown that DNNs, especially convolutional neural networks for image recognition (Krizhevsky et al., 2012; Simonyan and Zisserman, 2014; He et al., 2016), preserve the representational geometry in object-sensitive visual areas (Khaligh-Razavi and Kriegeskorte, 2014; Yamins et al., 2014; Cichy et al., 2016), and predicts neural and fMRI responses to natural picture or video stimuli (Güçlü and van Gerven, 2015b, a; Wen et al., 2016, 2017; Eickenberg et al., 2017; Seeliger et al., 2017), suggesting their close relevance to how the brain organizes and processes visual information (Cox and Dean, 2014; Kriegeskorte, 2015; Yamins and DiCarlo, 2016; Kietzmann et al., 2017; van Gerven, 2017). DNNs also open new opportunities for mapping the visual cortex, including the cortical hierarchy of spatial and temporal processing (Güçlü and van Gerven, 2015b, a; Cichy et al., 2016; Wen et al., 2016; Eickenberg et al., 2017), category representation and organization (Khaligh-Razavi and Kriegeskorte, 2014; Wen et al., 2017), visual-field maps (Wen et al., 2016; Eickenberg et al., 2017), all by using a single experimental paradigm with natural visual stimuli. It is even possible to use DNNs for decoding visual perception or imagery (Wen et al., 2016; Horikawa and Kamitani, 2017). Such mapping, encoding, and decoding capabilities all require a large amount of data from single subjects in order to train subject-specific models. Results in this study suggest that even 10 hours of fMRI data in response to diverse movie stimuli may still be insufficient for DNN-based encoding models (Fig. 2). Therefore, it is difficult to generalize the models established with data from few subjects to a large number of subjects or patients for a variety of potential applications.

The methods developed in this study fill this gap, allowing DNN-based encoding models to be trained for individual subjects without the need to collect substantial training data from them. As long as models have been already trained with a large amount of data from existing subjects or previous studies, such models can be utilized as the prior models for a new subject and be updated with additional data from this subject. Results in this study demonstrate that with prior models, encoding models can be trained with 16-min video-fMRI data from a single subject to reach comparable encoding performance as the models otherwise trained with over two hours of data but without utilizing any prior models (Fig. 3). Apparently, data acquisition for 16 minutes readily fit into the time constraint of most fMRI studies. With the method described herein, it is thus realistic to train encoding models to effectively map and characterize visual representations in many subjects or patients for basic or clinical neuroscience research. The future application to patients with various cortical visual impairments, e.g. facial aphasia, has the potential to provide new insights to such diseases and their progression.

The methods developed for transferring encoding models across subjects might also be usable to transfer such models across imaging studies with different spatial resolution. The fMRI data in this study are of relatively low resolution (3.5mm). Higher resolution about 1mm is readily attainable with fMRI in higher field strengths (e.g. 7T or above) (Goense et al., 2016). Functional images in different resolution reflect representations in different spatial scales. High-field and high-resolution fMRI that resolves representations in the level of cortical columns or layers is of particular interest (Yacoub et al., 2008; Goense et al., 2016); but prolonged fMRI scans in high-field face challenges, e.g. head motion and susceptibility artifacts as well as safety concern of RF power deposition. Transferring encoding models trained with 3-T fMRI data in lower resolution to 7-T fMRI data in higher resolution potentially enables higher throughput with limited datasets. Note that transferring the encoding models is not simply duplicating the models across subjects or studies. Instead, new data acquired from different subjects or with different resolution serve to reshape the prior models to fit the new information in specific subjects or representational scales. It is perhaps even conceivable to use the method in this study to transfer encoding models trained with fMRI data to those with neurophysiological responses observable with recordings of unit activity, local field potentials, and electrocorticograms. As such, it has the potential to compare and converge neural coding in different spatial and temporal scales. However, such a potential is speculative and awaits verification in future studies.

This study also supports an extendable strategy for training population-wide encoding models by collecting data from a large group of subjects. In most of the current imaging studies, different subjects undergo the same stimuli or tasks with the same experiment paradigm and the same acquisition protocol (Hodge et al., 2016). Such study design allows for more convenient group-level statistics, more generalizable findings, and easier comparison across individuals. However, if one has to collect substantial data from each subject, it is practical too expensive or unrealistic to do so for a large number of subjects. An alternative strategy is to design a study for a large number of subjects, but only collect imaging data from subjects undergoing partly different visual stimuli, e.g. watching different videos. For the population as a whole, data with a large and diverse set of stimuli become available. The methods described herein lay the technical foundation to combine the data across subjects for training population-wide encoding models. This strategy may be further complemented by also using a small set of stimuli (e.g. 16-min video stimuli) common for all subjects. Such stimuli can be used to functionally align the data from different subjects to account for individual differences (Fig. 6) (Haxby et al., 2011; Conroy et al., 2013; Guntupalli et al., 2016). It also provides comparable testing data to assess individual differences.

In addition, our methods allow population-wide encoding models to be trained incrementally. For a study that involves data acquisition from many subjects, data are larger and growing. It is perhaps an unfavorable strategy to analyze the population data only after data are available from all subjects. Not only is it inefficient, analyzing the population data as a whole requires substantial computing resources – a common challenge for “big data”. Using online learning (Fontenla-Romero et al., 2013), the methods described herein allows models to be trained and refined as data acquisition progresses. Researchers can examine the evolution of the trained models, and decide whether the models have converged to avoid further data acquisition. As population-wide encoding models become available, it is more desirable to use them as the prior models for training encoding models for specific subjects, or another population. Populationwide models are expected to be more generalizable than models trained from one or few subjects, making the prior models more valid and applicable for a wide group of subjects or patients.

This study focuses exclusively on natural vision. However, the methods developed are anticipated to serve well for more general purposes, including natural language processing, speech and hearing.

## Acknowledgement

The authors are thankful to Dr. Xiaohong Zhu, Dr. Byeong-Yeul Lee, Yizhen Zhang, and Kuan Han for constructive discussion. The research was supported by NIH R01MH104402 and Purdue University.

